# Hydrogen peroxide as regeneration-initiation signal that activates pERK to trigger planarian regeneration

**DOI:** 10.1101/712356

**Authors:** Vincent Jaenen, Susanna Fraguas, Karolien Bijnens, Mireia Vaca, Tom Artois, Rafael Romero, Karen Smeets, Francesc Cebrià

## Abstract

Despite the extensive research on molecular pathways controlling the process of regeneration in planarians and other regeneration models, little is known about the actual initiation signals necessary to induce regeneration. Previously the involvement of ROS, EGFR and MAPK/ERK has been demonstrated during planarian regeneration, however the exact interplay has not been yet described. By selectively interfering with major mediators in all three key parts (ROS, EGFR & MAPK/ERK), we were able to identify amputation/wound-induced ROS, and H_2_O_2_ specifically, as upstream cue in activating regeneration-initiation. In addition, our results demonstrate new relationships between regeneration related ROS production and MAPK/ERK activation at early regeneration stages, as well as the involvement of the EGFR-signaling pathway. In summary, our results suggest a new and more extensive signaling model with ROS, and H_2_O_2_, highlighted as upstream initiation-factor and its important functions in the downstream EGFR-MAPK/ERK pathway during planarian regeneration.

## Introduction

Regeneration is the fascinating phenomenon in which animals are able to repair and regrow lost or damaged tissues, structures and even the whole body^1^. At the cellular level, this includes proliferation, migration and differentiation, processes that need to be under a strict genetic and molecular control^2^. The capacity to regenerate missing or damaged parts is widely distributed across the animal kingdom^3^. The level of regeneration varies greatly among different species; from full body regeneration in invertebrates such as hydra and planarians, to specific organs and structures as for example limb regeneration in amphibians and heart regeneration in zebrafish^4-6^. Mammals such as humans, on the other hand, have very limited regenerative capabilities. Understanding how tissue development takes place in regenerative animals might provide fundamental knowledge for the further improvement of regenerative medicine.

In this context, freshwater planarians are a model with several attractive features: i) they can regenerate a whole animal from a tiny piece of their body, ii) a large part of their body consists of a population of adult pluripotent stem cells called neoblasts, with a similar transcriptional profile as compared to other invertebrate and vertebrate stem cells, and iii) they use conserved signalling pathways to regulate cell differentiation, patterning and morphogenesis. As such, Hedgehog and Wnt/ β-catenin signaling are required to re-establish the anteroposterior axis of the animal, the BMP pathway regulates the dorsoventral axis, and the EGFR pathway is needed for proper neoblast differentiation^7-9^. Although several studies have uncovered a pivotal role of these, and other, signaling pathways during planarian regeneration, little is known about what upstream signals activate them to trigger a regenerative answer after an amputation. Recently, Owlarn and colleagues reported that a planarian extracellular signal-regulated protein kinase (ERK) was strongly activated in a stem cell-independent manner, just minutes after amputation. By inhibiting protein synthesis using cycloheximide, they showed that ERK activation was triggered by injury signals that did not originate from newly synthesized proteins. Therefore, ERK activation by a still unknown factor, appears to be the most upstream initiator of planarian regeneration^10^. In recent years, many *in vitro* as well as *in vivo* studies put forward reactive oxygen species (ROS) as upstream signaling molecules of regeneration^11-13^. H_2_O_2_ for example, can cross cell membranes through aquaporin channels and gap-junctions and diffuse freely between the cells^14,15^. Similar to what happens in other physiological processes such as growth, inflammation and ageing, wound-induced ROS, and more specifically H_2_O_2_ with a rather long half-life, can play a regulating role in early wound responses and regeneration. In 2013, Love *et al.* demonstrated that amputation-induced ROS production is absolutely required for Xenopus tadpole tail regeneration^16^. Shortly after that, Gauron and colleagues showed a similar ROS production at the amputation site during fin regeneration in adult zebrafish and proved its role in blastema formation^17^. More recently, the Serras group demonstrated the presence of an oxidative burst just minutes after inducing regeneration of the wing imaginal disc in Drosophila^18^. Also in planarians, we observed that an amputation-induced ROS burst is necessary for proper stem cell differentiation and successful regeneration^13^.

Many studies already indicated an interplay between ROS and mitogen-activated protein kinases (MAPK) signaling pathways^18-22^. Ruffels *et al*. simulated a ROS burst by direct exposure of exogenous H_2_O_2_ to human neuroblastoma cells, resulting in a significant increase of ERK activation. In Drosophila, ROS-dependent stimulation of MAPKs is essential for the activation of JAK/STAT signaling, which drives regeneration. In addition, literature shows that the activation of MAPK pathways can be mediated by the upstream epidermal growth factor receptor (EGFR) in planarians^24,25^ and other models^26-29^. Also, the activation of the EGFR pathway by ROS has been described in different models^30,31^. Furthermore, previous studies have identified a functional relationship between the EGFR pathway and the egr (early growth response) family of transcription factors in planarian regeneration, in which a link between egr genes and ERK activity has been demonstrated in different models^32-35^. Overall, these reports suggest that wound-induced ROS signaling operates through EGFR-MAPK pathways in order to stimulate transcriptional expression of cytokines that in turn will be crucial to trigger tissue repair and restore homeostasis^19^.

Even though ROS, EGFR and ERK have been shown to be required for planarian regeneration, it is not known the relationship between these pathways. Here, we show that amputation-induced ROS production might be the upstream cue that would activate ERK signaling to initiate regeneration in these animals.

## Results

### Generation of Reactive Oxygen Species after applying an R - or H wound

Recently, it was shown that a common generic wound response program is triggered by both injuries that require only wound healing (H-wounds) and by injuries that imply tissue loss and, therefore, require regeneration (R-wounds)^10^. In previous research, we showed an amputation-induced ROS burst after inflicting an R-wound in planarians^13^. Here, we confirm the fast ROS production at the R-wound site (Fig. **1**A.1), and additionally show that a ROS burst occurs after applying an H-wound (Fig. **1**A.2). Negative controls (without carboxy-H2DCFDA), showed no autofluorescence at both R- and H-wound sites (Suppl. Fig. **1**).

**Figure 1.**
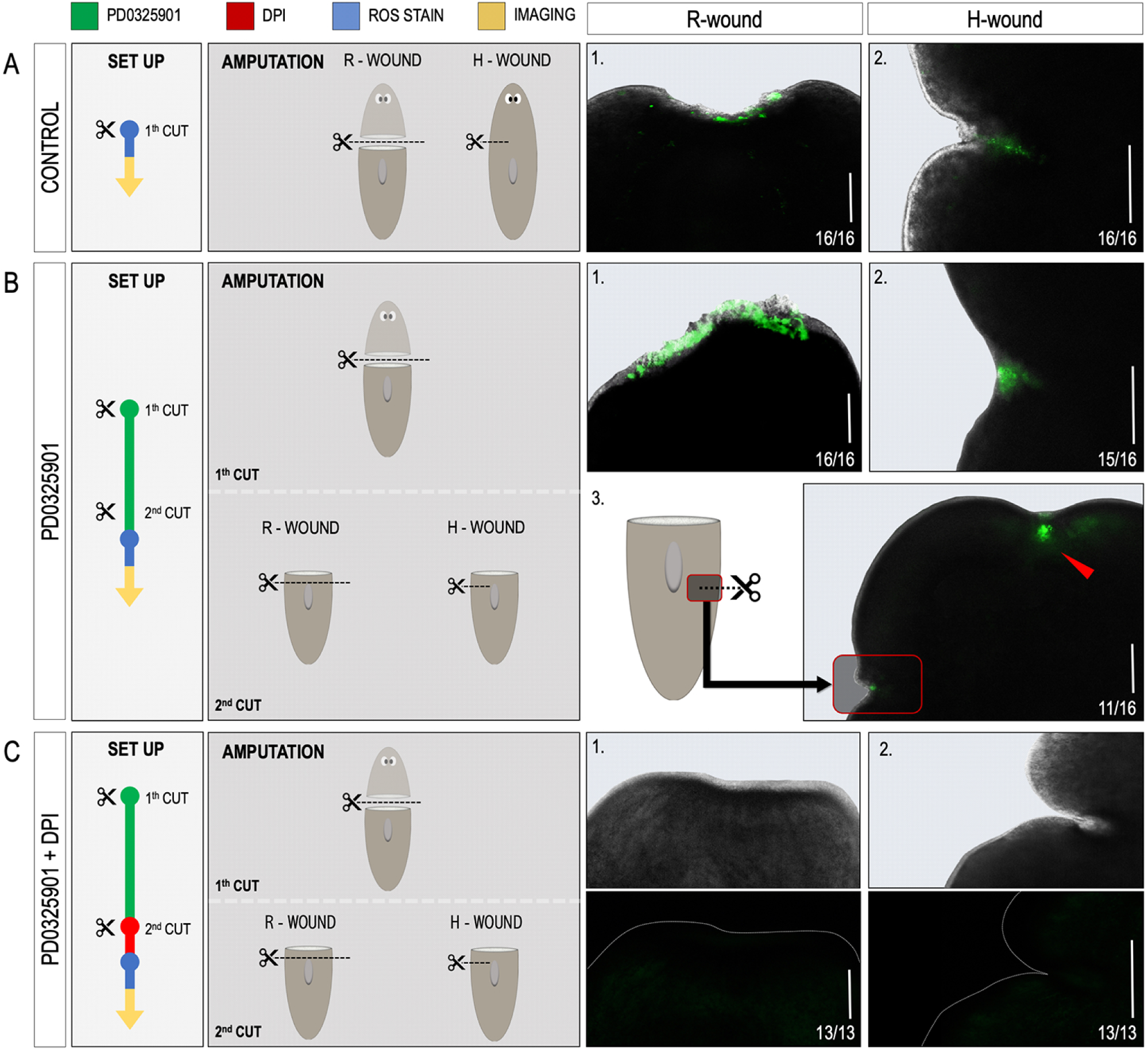
*In vivo* visualisation of ROS production at the amputation site after re-wounding MEK-inhibited fragments. For each condition, the experimental setup is displayed in the white boxes. Colour code as follows; green: MEK inhibition by PD0325901 (10 µM, 5 days), red: inhibition of ROS production by DPI (3 µM, 5 hours), blue: ROS visualisation procedure using carboxy-H_2_DCFDA, yellow: imaging procedure. The amputation setup is displayed in the dark grey boxes. Regenerative wound (R-wound) & healing wound (H-wound). All animals were visualised 30 minutes post amputation (MPA). A representative close-up merged image of bright field and fluorescence of either R-wounds or H-wounds is displayed on the right panel. (**A**) ROS production visualisation at the site of an R-wound (**A.1**) and an H-wound (**A.2**) in controls. (**B**) ROS production visualisation at the amputation site of an R-wound (**B.1**) and an H-wound (**B.2**) in MEK-inhibited (PD0325901), “dormant” fragments. (**B.3**) Image of the site of the original R-wound applied before MEK-inhibition (red arrowhead) together with the newly applied H-wound (red square). (**C**) ROS production visualisation after rewounding (**C.1**: R-wound, **C.2**: H-wound) MEK-inhibited and DPI exposed dormant fragments. Because of the strongly reduced fluorescence in (**C**), the close-up of the wound site is shown in bright field (upper panels) and fluorescence (lower panels) separated. The dotted, white line indicates the border of the wound site. Scale bars 100µm.

Owlarn and colleagues have demonstrated the triggering of generic initiation-signals in both wound types in dormant, MEK-inhibited fragments^10^. Therefore, it is important to verify the presence of ROS at the wound site after rewounding MEK-inhibited fragments in order to analyze its possible role in the rescue of regeneration. ROS were indeed detected at the site of the newly inflicted R-wound (Fig. **1**B.1, 16/16) as well as H-wound (Fig. **1**B.2, 16/16) in these MEK-inhibited fragments. Interestingly, ROS levels were not only increased at the wound site after inflicting an H-wound, (Fig. **1**B.3, red square) but also occurred at the original, dormant, R-wound sites (Fig.**1**B.3, red arrow head, 11/16). To confirm the ROS signal, DPI was used as a negative control (Fig. **1**C, 13/13).

In summary, these results, together with the observed regenerative impairments after ROS inhibition (Suppl. Fig. **2**)^13^, point out the fast developed, wound-induced ROS as putative upstream key mediator in both initiating regeneration in control animals and in the rescue of regeneration in dormant MEK-inhibited fragments.

### Hydrogen peroxide treatment rescues regeneration in dormant MEK-inhibited fragments

To functionally confirm a role for ROS as a regeneration initiating signal, we investigated if we could reactivate MEK-inhibited, dormant blastemas by treating them with an exogenous ROS-source. The *in vivo* ROS stain that we addressed to visualise wound-induced ROS production detects a broad range of reactive oxygen species, making it difficult to specify which ROS to use. However, because of the beneficial characteristics of H_2_O_2_ for its function as messenger molecule and the supporting evidence regarding the activation potency for several molecular pathways, we focused on using H_2_O_2_ as exogenous ROS-source (Fig. **2**).

**Figure 2.**
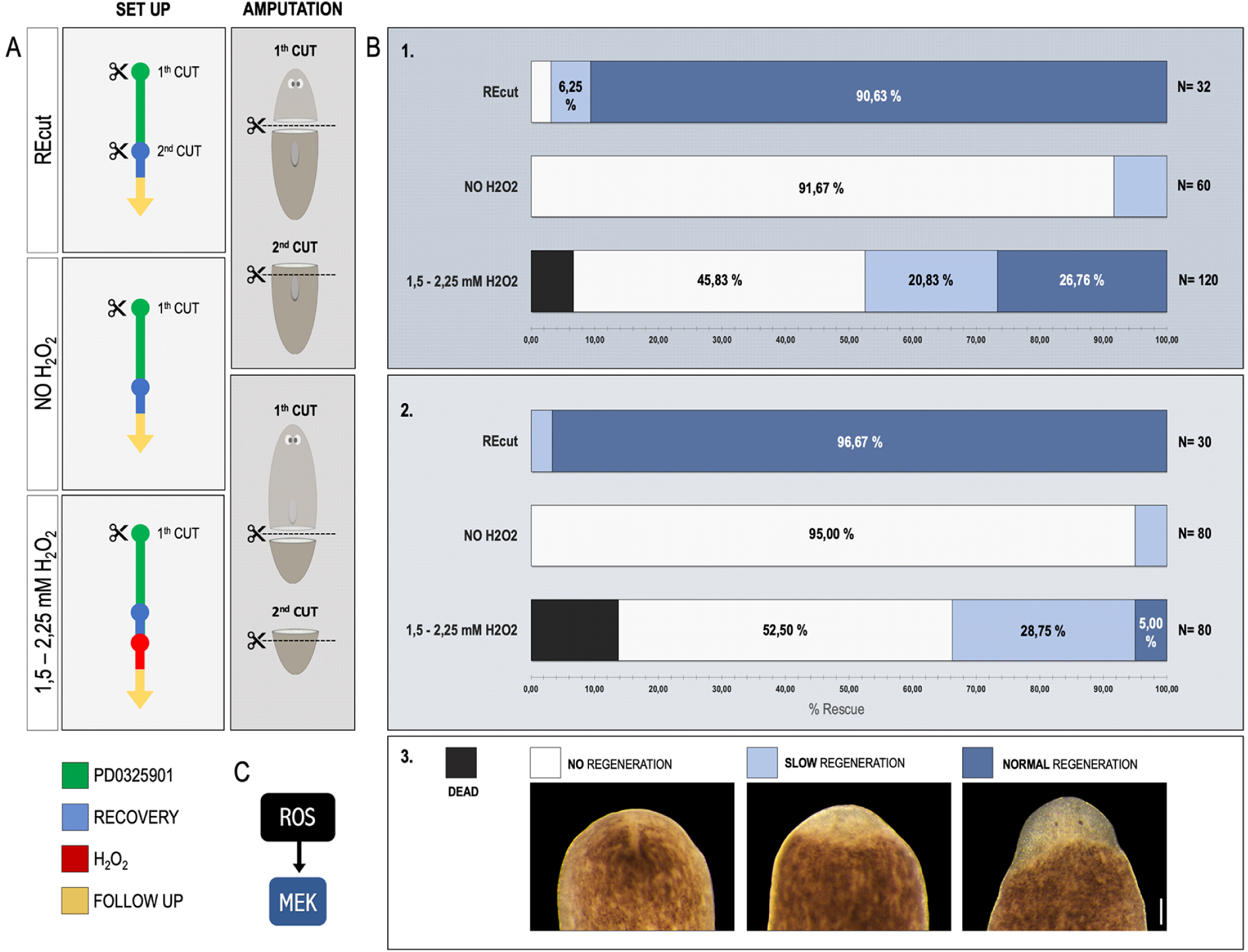
H_2_O_2_ treatment rescues regeneration in MEK-inhibited dormant fragments. (**A**) For each condition, the experimental setup is displayed in the white boxes. Colour code as follows; green: MEK inhibition by PD0325901 (10 µM,5 days), blue: recovery in fresh medium after several washes (1day), red: treatment with H_2_O_2_ (1,5-2,25 mM, 6 hours), yellow: follow up in fresh medium after several washes (7 days). The amputation setup is displayed in the dark grey boxes; 1^st^ cut applied just before MEK-inhibition, 2^nd^ cut applied after recovery period and only applicable in the “REcut” condition). (**B**) “Dormant”, MEK-inhibited, trunk fragments with pharynx (**B.1**) and tail fragments without pharynx (**B.2**) were Recut (2^nd^ cut) or treated with H_2_O_2_. “NO H_2_O_2_” controls were neither recut or treated with H_2_O_2_, and were only MEK-inhibited. (**B.3**) Display of the different regenerative outcomes (7 days post rewounding or H_2_O_2_ treatment) quantified on graphs B.1 and B.2: no regeneration (white), slow regeneration (light blue), normal regeneration (dark blue) and dead (black). “Normal regeneration” is considered as full regeneration including eyes, and if needed pharynx. In the case of “slow regeneration”, a clear blastema was visible, however, no eyes and/or pharynx were differentiated at this time point. “No regeneration” refers to the absence of a blastema and a total block of regeneration. Sample numbers are indicated on the right site of each graph. (**C**) Putative model in which ROS would function upstream of MEK.Scale bars 100µm.

When inflicting a new R-wound (Fig. **2**B: REcut) to MEK-inhibited trunk fragments (with pharynx), 96,88% (n=32) of the fragments regenerated. A small fraction of these animals (6,25%) started regenerating, but neither eyes nor pharynx could be identified at the moment of comparison. In tail fragments (without a pharynx), 100% (n=30) restarted regeneration after inflicting an R-wound, of which 96,67% regenerated normally. In the absence of a new R-wound (Fig. **2**B: NO H_**2**_O_**2**_), 91,67% (n = 60) of the MEK-inhibited trunks failed to regenerate, while the remaining part of the worms regenerated slower. The same trend was visible in the tail fragments: 95% (n=80) of the fragments did not regenerate and in 5% regeneration was delayed. In all cases, the absence of regeneration remained for at least 21 days. Initial dose-response experiments indicated that a 6 hours treatment with 1,5 to 2,25mM H_**2**_O_**2**_ was enough to fully rescue 26,76% (n=120) of the trunk fragments (with a pharynx), and 5% (n=80) of tail fragments (without a pharynx). An additional 20,83% of the trunk fragments were partially rescued, showing a slower regeneration. In case of the tail fragments this partial rescue reached 28,75%. In both setups, trunk and tail fragments, a small amount of animals died; 6,58% and 13,75% respectively. In order to exclude wounding by H_2_O_2_, a ROS staining was performed on dormant fragments after 3 and 6 hours of exposure to H_2_O_2_. No ROS-induced wounding was observed at the epidermis of the H_2_O_2_-treated planarians (Suppl. Fig. **3**).

Taken together, these results show the potency of H_2_O_2_ in reversing MEK-inhibition, consequently rescuing regeneration. Hereby, it supports the finding of ROS as upstream regeneration-initiation signal.

### Inhibition of ROS production blocks ERK activation at the wound site

To identify downstream targets and a possible interaction between the amputation-induced ROS burst and ERK, we performed an immunostaining with an anti-pERK antibody after ROS inhibition with 3 µM DPI^23 24^. In the control conditions, where animals were kept in regular medium or DMSO, an activation of pERK was observed at the wound site of head- (control: 4/5, DMSO: 3/6), trunk- (control: 6/7, DMSO: 6/8) as well as tail pieces (control: 4/6, DMSO: 4/6). This pERK-sinal was strongly reduced in the DPI-treated fragments (heads: 5/7, trunks: 7/9, tails: 5/7) (Fig. **3**B), suggesting that an amputation-induced ROS production at the wound site is required for the phosphorylation and proper activation of ERK (Fig. **3**C).

**Figure 3.**
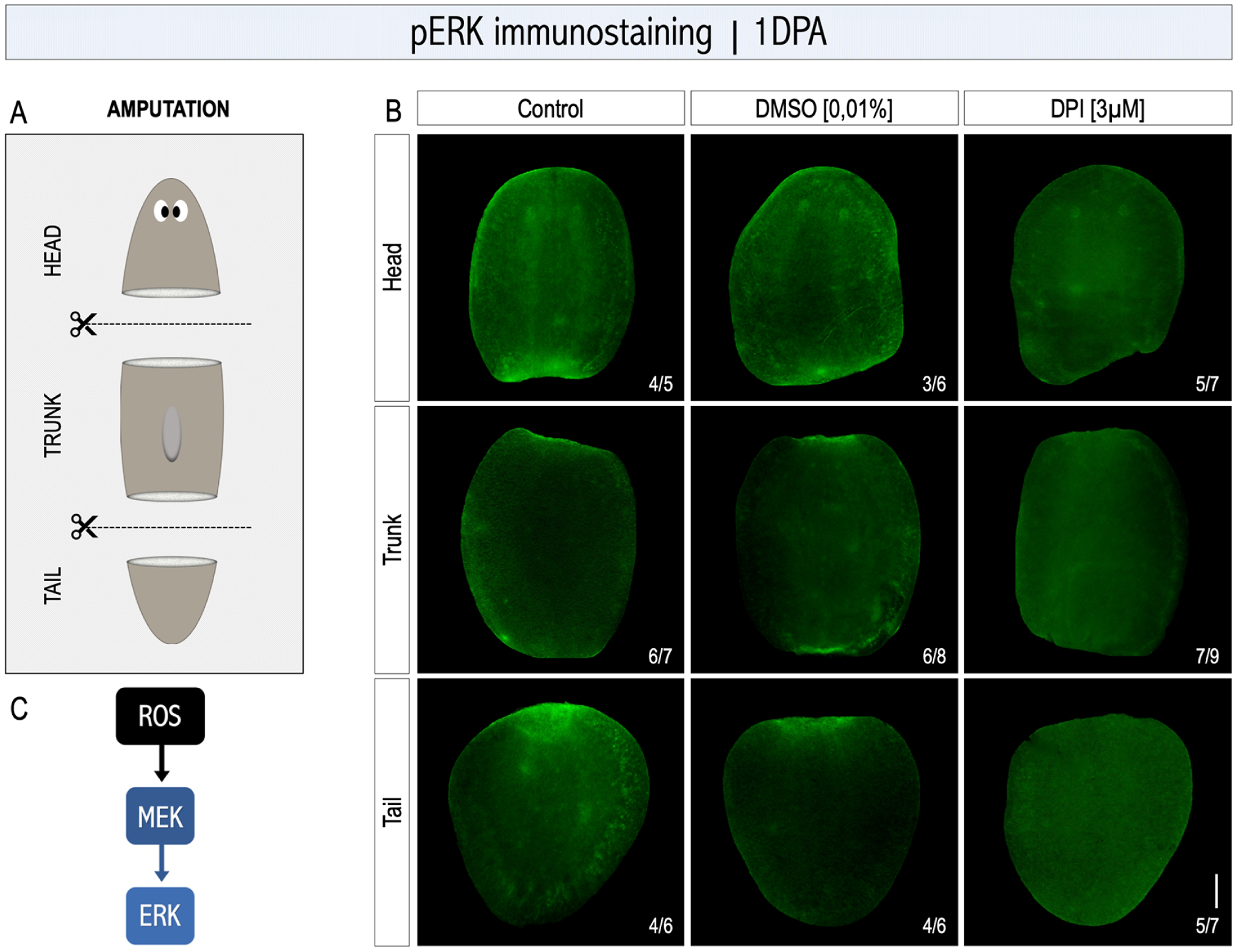
pERK activation decreases after the inhibition of ROS production. (**A**) The amputation setup is displayed in the grey box, indicating the amputation sites. (**B**) Immunostaining with an anti-pERK antibody on 1-day regenerating fragments. Animals kept in culture medium or 0,01% DMSO were used as control animals, while in the treatment group, ROS production was inhibited with DPI (3 µM) administered in the culture medium. All pictures were taken under the same light intensity settings. Sample numbers are indicated in the images. (**C**) Putative model in which ROS could activate pERK through MEK. Scale bars 100µm.

### *Smed-egfr-3* and *Smed-egr-4* silencing impair ROS production and pERK activation in regenerating animals

Literature has already described examples of the activation of EGFR signaling by ROS and the potential of EGFR in mediating MAPK signaling including ERK activation^24-31^. Additionally, the egr (early growth response) family of transcription factors is suggested to be a downstream target of EGFR signaling in planarian regeneration as well as in other models^32-35^.

In order to further investigate the functional relationship of *Smed-egfr-3* and *Smed-egr-4* with pERK, we carried out an immunostaining with the anti-pERK antibody in controls and RNAi-mediated *Smed-egfr-3* and *Smed-egr-4* knockdown animals. (Fig. **4**A). A clear pERK activation was observed in all control fragments at 6HPA (trunk pieces: 7/11, tail pieces: 7/8) and 1DPA (trunk pieces:8/9, tail pieces: 11/14). In contrast, a strong reduction of the anti-pERK signal at the wound site was observed in *Smed-egfr-3* silenced fragments at 6HPA (trunk pieces: 5/5, tail pieces: 5/7) and 1DPA (trunk pieces:7/8, tail pieces:12/14). Similar results were observed after silencing *Smed-egr-4*, both at 6HPA (trunk pieces: 9/9, tail pieces: 7/7) and 1DPA (trunk pieces:6/7, tail pieces:8/9) (Fig. **4**B). These results suggest that *Smed-egfr-3* and *Smed-egr-4* are required for ERK activation (Fig. **4**C).

**Figure 4.**
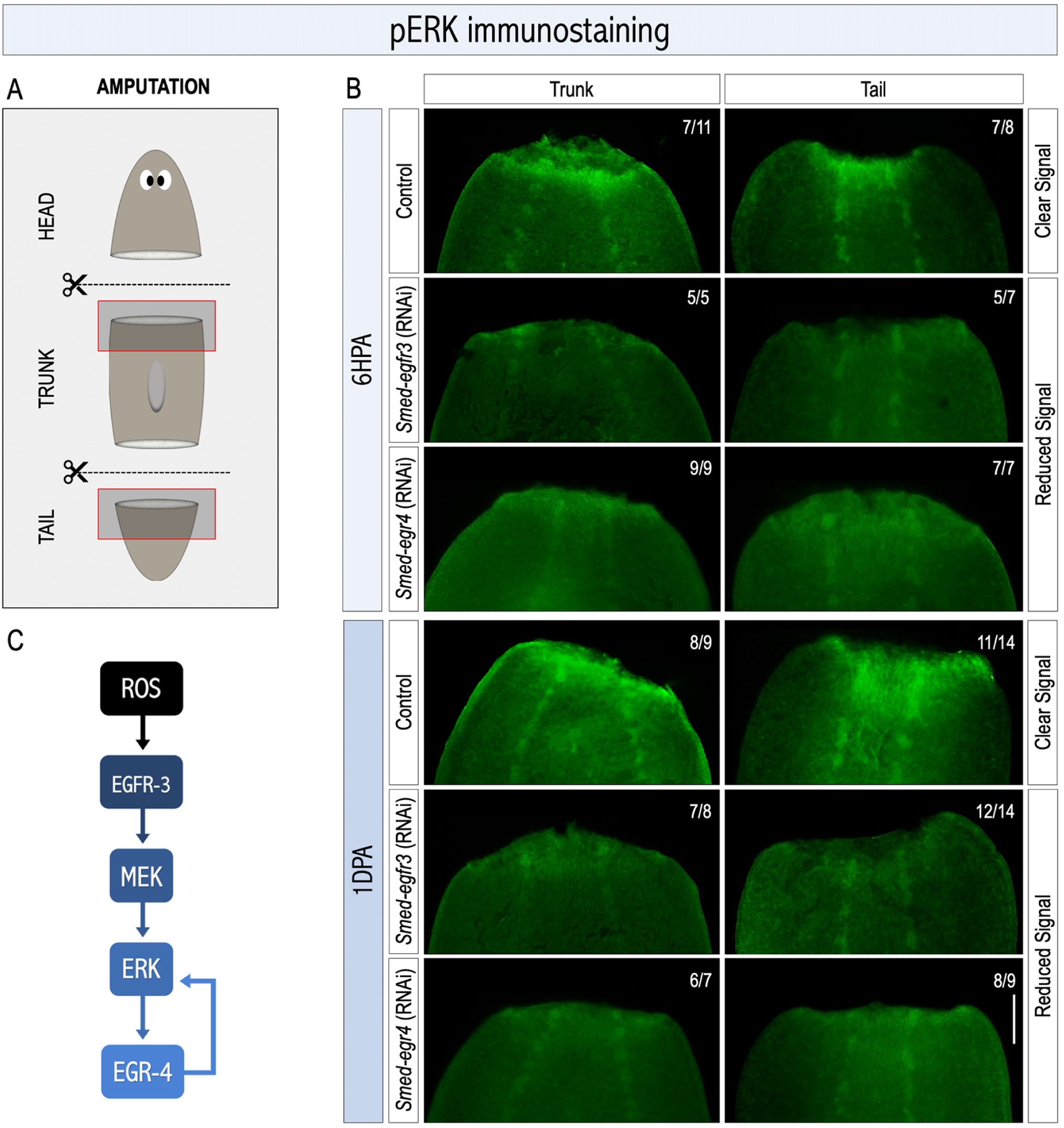
pERK activation regulation by *Smed-egfr-3* and*Smed-egr-4*. (**A**) The amputation setup is displayed in the grey box, indicating the amputation sites. Red squares correspond to the anterior blastemas shown in B at 6 hours and 1 day post-amputation. (**B**) Immunostaining with an anti-pERK antibody in controls, after *Smed-egfr-3* or *Smed-egr-4* RNAi. All pictures were taken under the same light intensity settings. In all panels anterior towards the top. Sample numbers are indicated in the images. (**C**) Putative model in which ERK activation might be regulated by *Smed-egfr-3* and *Smed-egr-4*. Scale bars 100µm.

To characterize the link of ROS with EGFR signaling during planarian regeneration, an *In vivo* ROS visualisation was performed in controls and animals subjected to RNAi silencing of either *Smed-egfr-3* or *Smed-egr-4*. (Fig. **5**). Whereas the presence of the amputation-induced ROS burst was clear in the control animals (30MPA: 5/7, 6HPA: 7/8, 24HPA: 6/8), trunk fragments subjected to *Smed-egfr-3* knockdown showed a clearly diminished ROS production at all time points (30MPA: 6/7, 6HPA: 5/7, 24HPA: 5/7). Similar results were observed in *Smed-egr-4* RNAi animals (30MPA: 6/9, 6HPA: 4/7, 24HPA: 5/8) (Fig. **5**B).

**Figure 5.**
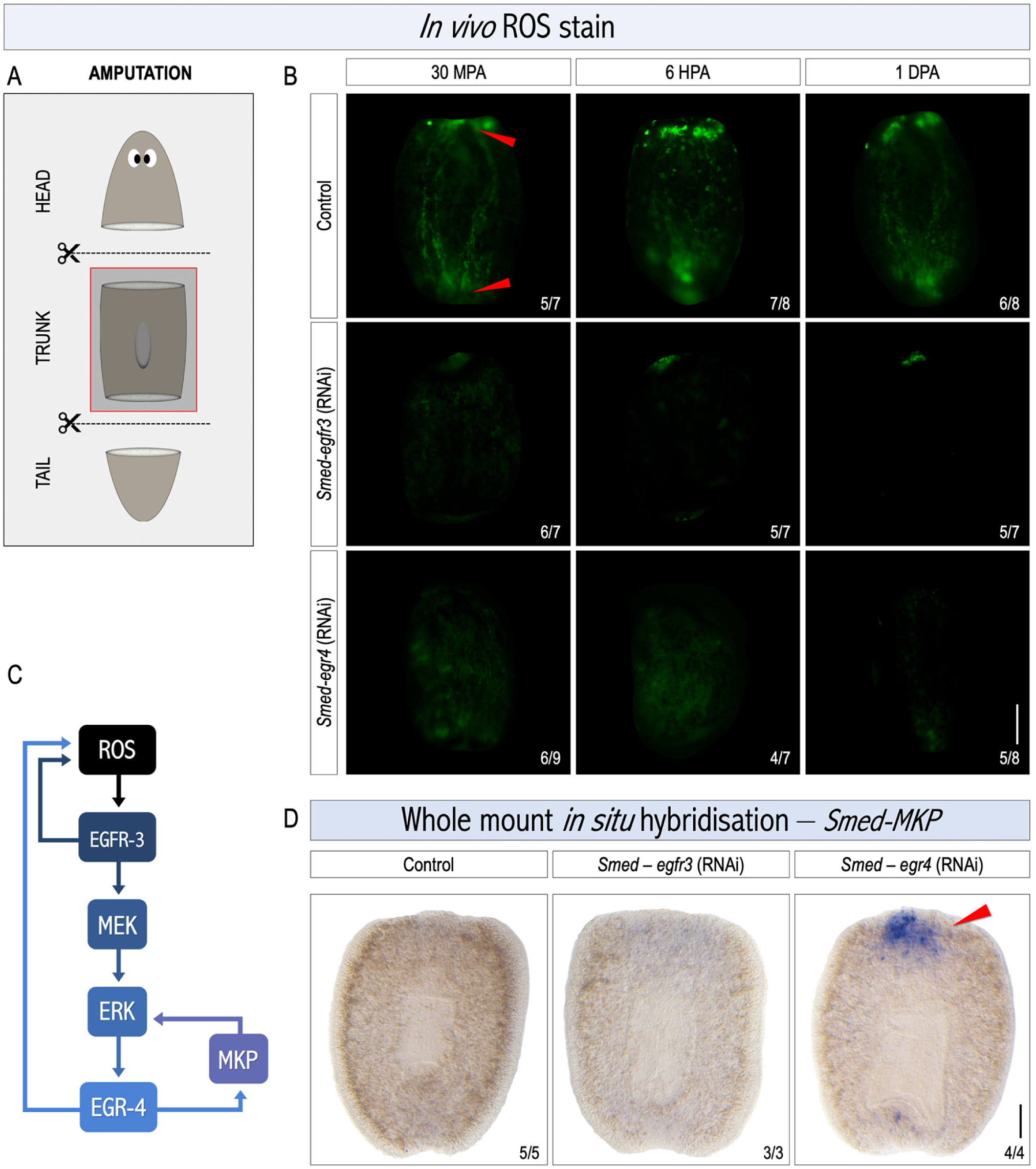
ROS production is regulated by *Smed-egfr-3* and *Smed-egr-4* during early regeneration. (**A**) The amputation setup is displayed in the grey box, indicating the amputation sites. The red square corresponds to the images shown in (**B**) and (**D**). (**B**) *In vivo* visualisation of amputation-induced ROS levels 30 minutes -, 6 hours - and 1 day post amputation (MPA, HPA, DPA) in controls and *Smed-egfr-3* or *Smed-egr-4* RNAi fragments. All pictures were taken under the same light intensity settings. Sample numbers are indicated each image. Scale bar 100µm. (**C**) Putative model showing the possible relationships between ROS, EGFR-3, ERK, egr-4 and mkp. (**D**) Whole-mount in situ hybridization of *Smed-mkp* in controls and *Smed-egfr-3* or *Smed-egr-4* RNAi planarians, 6 hours post-amputation. All panels are oriented with the anterior towards the top. Sample numbers are indicated each image. Scale bar 200µm.

Altogether, these results indicate that *Smed-egfr-3* as well as *Smed-egr-4* might play a pivotal role in regulating the amputation-induced ROS production, possibly through the existence of feedback mechanisms (Fig. **5**C). Furthermore, it is suggested that the activation of pERK by ROS could be mediated by the EGFR pathway during regeneration. Additionally, results point out the existence of a second, negative feedback mechanism of egr-4 in order to regulate pERK activation as well (Fig. **4**C).

### MAPK phosphatase is overexpressed in *Smed-egr-4* knockdown animals

In the context of previously suggested negative feedback mechanism of egr-4 in relation to ERKs activation, a study of Tasaki *et al.* 2011 showed that in planarians pERK could be negatively regulated by the action of a MAPK phosphatase (MKP)^23^. Therefore, the expression of *Smed-mkp* was checked in controls and planarians subjected to knockdown of either *Smed-egfr-3* or *Smed-egr-4*. A strongly increase of *Smed-mkp* expression was observed at the anterior wound site of the regenerating trunk fragments in *Smed-egr-4* RNAi animals (n=4/4) in comparison to no expression in both control (n = 5/5) and *Smed-egfr-3* RNAi (n = 3/3) animals (Fig. **5**D). These results suggest that *egr-4* might be required to maintain pERK activation by inhibiting *Smed-mkp*.

## Discussion

The complex process of animal regeneration is highly organized and regulated by a tightly controlled network of signaling pathways, of which we have only discovered the tip of the iceberg. Till now, regeneration research has mainly focused on molecular mechanisms regulating cell fate, polarity re-establishment, tissue differentiation and organ positioning, while the question on what initially triggers regeneration remains unanswered in many cases. In recent years several studies have reported the indispensability of reactive oxygen species (ROS) during regeneration. Amputation-induced ROS production was observed in early tail regeneration of Xenopus tadpoles and in fin regeneration of adult zebrafish^16,17^. Similar observations were reported during the regeneration process of the wing imaginal disc in Drosophila and during anterior and posterior regeneration of the planarian *Schmidtea mediterranea*^13,18^. In all of the above-mentioned cases, ROS production was clearly linked to the capacity to regenerate, e.g. the absence of ROS production during regeneration-initiation led to regenerative impairments (Suppl. Fig. 1)^10,13,16-18^.

Previously we showed a rapid ROS burst at the wound site after inflicting a regenerative(R)-wound as well as its necessity for proper regeneration in planarians^13^. Here, we demonstrate that ROS are also produced after healing(H)-wounding (Fig. 1A). Owlarn and colleagues showed that in planarians both R-wounds and H-wounds trigger a common initial molecular response mediated by ERK signaling. Inhibition of the ERK upstream activator, MAPK/ERK kinase (MEK), completely blocks regeneration after an R-wound^10^. No blastema is initiated and the body fragments remain “dormant” until a new R-wound or H-wound is made. The fact that inflicting a new H-wound is capable of rescuing regeneration of dormant fragments suggests that, in planarians, any kind of wounding (whether or not it results in tissue loss) triggers an early response through ERK activation and only in the context of tissue loss this is followed by a late regenerative response that leads to the restoration of the missing part(s)^10^. However, the initial upstream signal that activates ERK, remained unknown. Here, we show a ROS burst in the MEK-inhibited dormant fragments after inflicting a new H- or R-wound (Fig. 1B). Surprisingly, H-wounding not only induces a ROS burst at the H-wound site but also at the original, dormant R-wound site (Fig. 1B3), outlining again the involvement of ROS in the initiation of regeneration. However, in planarians it is not yet clear if ROS can propagate as signaling molecules by themselves or if they induce an unknown secondary signal to the dormant wound site in order to locally restart regeneration^14,15,19^.

Generally it is stated that ROS function as messenger molecules and regulate processes such as cell proliferation, differentiation and patterning by modulating gene transcription and protein phosphorylation^11,12,19,36-39^. In many of these pathways, MAPKs are predominantly mentioned as a required intermediate link, playing pivotal roles in signal transduction from the cell membrane to the nucleus. Exogenous addition of H_2_O_2_ or treatment with ROS-inducing compounds, lead to the activation of the MAPK pathway^19,40,41^. On the other hand, the inhibition of ROS production or the stimulation of antioxidant defense mechanisms, block MAPK activation^20,42^. Taking into account that, in planarians, ROS are necessary for regeneration, that ROS are induced very early after H- and R-wounding, and that ERK activation is required for regeneration, we searched to determine whether ROS act upstream of ERK and which factors mediate ROS production and ERK activation. Our results show that MEK-inhibited dormant tails can be rescued by the addition of exogenous H_2_O_2_ without the need to inflict a new wound (Fig. 2). It is important to point out that the H_2_O_2_ treatment did not induce wounds in the epidermis, indicating that the rescue was not introduced by re-wounding the dormant tails (Suppl. Fig.3). We hypothesize that because of its relatively long half-life and good membrane permeability, H_2_O_2_ functions as secondary messenger and triggers the activation of important regeneration-related downstream signaling processes such as ERK activation. The strong reduction of ERK activation at the wound site after inhibition of ROS production in regenerating animals, again verifies the upstream function of ROS relative to ERK (Fig. 3).

In many organisms, the epidermal growth factor receptors (EGFR), one of families of the receptor tyrosine kinases (RTK), are known to regulate several biological processes by activating key downstream pathways including the MAPK pathway^26-29,43-45^. In planarians, *Smed-egfr-3* is required for proper regeneration as well as for ERK activation (Fig.4)^24,46^. Recent literature suggested that RTK-associated activation mechanisms are under redox control. Activation of EGFR signaling by ROS can occur in at several ways^47^. On one hand, intracellular ROS can facilitate the phosphorylation of EGFR and induce a subsequent cascade of phosphorylations. According to Peus and colleagues, H_2_O_2_ specifically acts as a critical mediator in this case^48^. The absence of ERK activation after inhibition of ROS production (Fig.3), and the rescue of regeneration in dormant tails by H_2_O_2_ in our results support this hypothesis (Fig.2). Furthermore, silencing *Smed-egfr-3* results in decreased amputation-induced ROS production, suggesting the dependency of ROS production on the EGFR signaling (Fig.5). This could be explained by the fact that ligand-dependent dimerization of EGFR induces ROS production for its autophosphorylation and consequent activation, leading to a decreased ROS production when silenced^47^. However, literature also suggests that H_2_O_2_ acts as a critical mediator in the ligand-independent phosphorylation and activation of EGFR^48^. Additionally, EGFR signaling can also be redox-controlled via MAPK phosphatases (MKPs), responsible for the dephosphorylation and inactivation of ERK. MPKs contain catalytic cysteine residues which are targets for oxidation by ROS leading to its deactivation^19,49^. As a result, ROS inhibition would lead to increased MKP activity and therefore inhibition of ERK activation.

Our results suggest a second regulator of MKP activity, namely *Smed-egr-4. Smed-egr-4*, codes for a zinc finger transcription factor of the early growth response factor family and is reported as a putative target of *Smed-egfr-3*^32^. The transcription factor is upregulated immediately after inflicting an R- or H-wound, while hardly expressed in intact planarians^50,51^. *Smed-egr-4* silencing results in regeneration defects similar to those after *Smed-egfr-3* downregulation, inhibition of ERKs activation and decreased ROS production^13,23,32,46^. Silencing of *Smed-egr-4* leads to a strong reduction of ERK activation together with a pronounced upregulation of *Smed-mkp* (Fig. 5) which suggests that *Smed-egr-4* might function as an inhibitor of MKP activity, preventing the dephosphorylation and inactivation of ERK. Together with ROS they form a secure and self-reinforcing regulation mechanism of MKP activation state. Moreover, the silencing of *Smed-egr-4* also leads to a reduced amputation-induced ROS production which suggest the existence of a feedback mechanism regulating ROS production by *Smed-egr-4* (Fig. 4). Because regulation of NADPH-oxidases (Nox) expression can be controlled by ERK-activation and linked transcription factors, *Smed-egr-4* knockdown which leads to increased MKP activation and consequently decreased ERK activation, can alter Nox-expression, explaining the impaired ROS production^53^. However, further experiments are necessary to clarify the involvement of Nox genes in planarians. All these data together with what is known about ROS and EGFR-MAPK signaling in other systems allows us to propose a model for the interactions of all these elements during planarian regeneration (Fig. 6).

**Figure 6.**
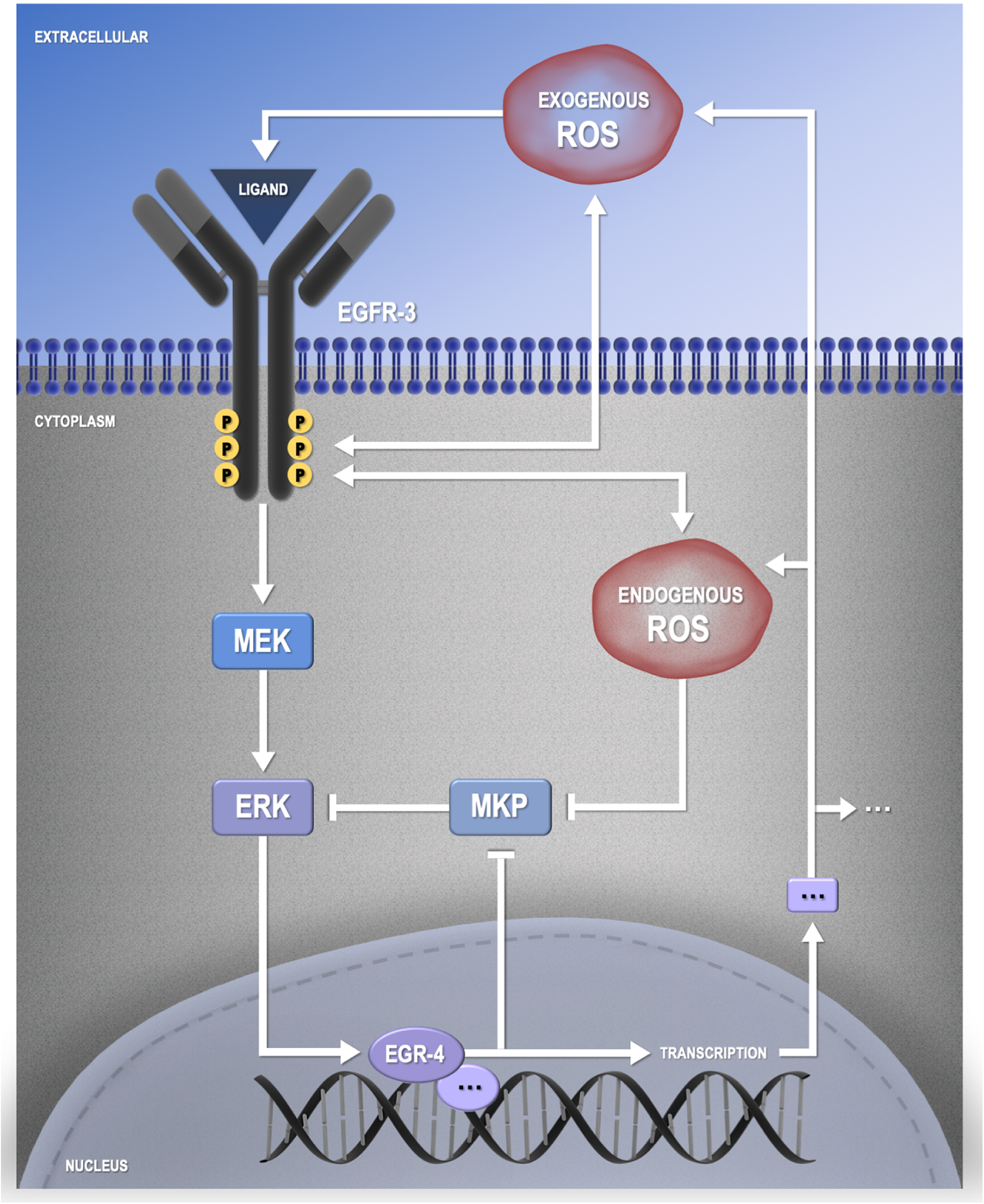
Proposed signaling model of interactions between ROS and the EGFR/MAPK pathway to regulate planarian regeneration. A possible regulatory pathway implicating ROS, egfr-3, egr-4 and MKP is displayed based on literature and the results presented here..

In summary, our results suggest that: i) ROS have the potential to rescue regeneration in MEK-inhibited dormant tails, ii) ROS are required for ERK activation at early regeneration stages, iii) the EGFR pathway can mediate ROS production with ERK activation during planarian regeneration. We provide the first evidence of amputation- and wound-induced ROS production in relationship with the EGFR-MAPK signaling pathway during planarian regeneration in which ROS not only are identified as most upstream trigger for regeneration-initiation, but additionally also perform its functions more downstream.

## Materials and Methods

### Planarian cultivation

An asexual strain of the freshwater planarian species *Schmidtea mediterranea* was kept in Milli-Q water containing 1.2 mM NaHCO_**3**_, 1.6 mM NaCl, 1.0 mM CaCl_**2**_, 1.0 mM MgSO_**4**_, 0.1 mM MgCl_**2**_ and 0.1 mM KCl (cultivation medium). The planarians were continuously maintained in the dark at a temperature of 20°C. Once a week they were fed with veal liver. Animals used in experiments were starved for at least 7 days before the procedure.

### Inhibition of Reactive Oxygen Species (ROS) Production

The nonspecific flavoprotein inhibitor, Diphenyleneiodonium chloride (DPI, Sigma Aldrich, D2926), was used in order to block ROS production by interfering with several electron transporters. Animals were exposed to 3µM DPI for 5 hours prior to *in vivo* ROS staining and 1hour prior amputation when followed by pERK immunohistochemistry. In both cases, animals were continuously exposed to DPI during the whole regeneration period. Because of its hydrophobic character, DPI was prepared in 0.01% dimethylsulfoxide (DMSO, Sigma Aldrich, 471267). In all experiments concerning DPI exposure, a DMSO-exposed control group was added to take into account the possible effects of DMSO since relatively high concentrations can have neurotoxic effects and influence cell proliferation in *S. mediterranea*^52^.

### Immunohistochemistry

To analyse the relationships between ROS, *Smed-egfr-3* and *Smed-egr-4* relative to ERK activation in early regeneration, a pERK immunostaining was performed after interfering with the aforesaid. Planarians were amputated followed by 6 and/or 24 hours of regeneration. Next, they were fixed and processed as previously described by Fraguas *et al.*^24^. Bleached animals were washed with PBSTx (1x PBS (10x PBS: 1,37M NaCl, 27mM KCl, 100mM Na_2_HPO_4_, 20mM KH_2_PO_4_in ultrapure H_2_O) and incubated for 4 hours in 1% blocking solution (1% BSA in PBSTx) followed by the primary antibody (anti-pERK23 diluted 1/1000 in blocking solution) overnight at 4°C. After PBSTx washes and 1 hour in blocking solution, they were incubated with the secondary antibody (goat anti-rabbit-POD diluted 1/500 in blocking solution) overnight at 4°C. After PBSTx washes, samples were incubated for 8 minutes in TSA Plus Fluorescein solution (1/50 TSA Plus Fluorescein in 1x Amplification Buffer (Tyramide Signal Amplification Labeling Kit No. 2; Molecular Probes, Thermo Fisher Scientific)) in darkness. Samples were mounted after the final PBSTx washes (RT) and analysed with a MZ16F fluorescence stereomicroscope (Leica) equipped with a ProgRes C3 camera (Jenoptik, Jena, Germany).

### MEK Inhibition

The chemical compound PD0325901 (Calbiochem) was used to reversibly inhibit MEK activity and subsequently prevent the phosphorylation and activation of ERK. As a consequence we obtained dormant planarian fragments as described by Owlarn *et al*.^10^. PD0325901 was dissolved in DMSO and used in a concentration of 25 µM. Planarians were exposed to PD0325901 for 1 hour prior and up to 5 to 7 days post amputation. The exposure solution was replaced daily or every two days, depending on the experiment. After treatment with PD0325901, animals were gently washed and placed into fresh medium until used for experiments.

### H_2_O_2_Treatment

Dormant, MEK-inhibited fragments were exposed to H_2_O_2_ with the intention to rescue regeneration. After initial range finding experiments, dormant fragments were exposed to either 1,5mM (0,005%) or 2,25mM (0,0075%) H_2_O_2_ (in cultivation medium) for 6 hours. After 3 washes with cultivation medium, they were kept in fresh medium.

### Reactive Oxygen Species (ROS) Detection

The compound 5-(and-6)-carboxy-2’,7’-dichlorodihydrofluorescein diacetate (carboxy-H_2_DCFDA, Image-iT LIVE Green Reactive Oxygen Species Detection Kit, Molecular Probes; Invitrogen, I36007) was used to visualise the *in vivo* production of ROS, in which fluorescent carboxy-DCF is produced through ROS oxidation. The ROS visualisation procedure was performed on *Smed-egfr-3* and *Smed-egr-4* RNAi knockdown animals as well as control – and MEK-inhibited animals either combined or not with the inhibition of ROS production by DPI. Animals were exposed to carboxy-H_2_DCFDA (25µM, 1ml) for 1 hour prior to amputation and for 1 day post RNAi. Amputated animals were again incubated in carboxy-H_2_DCFDA for 15 minutes before immobilisation in 2% low melting point (LMP) agarose (Invitrogen, 16520-050). Imaging of the samples was performed on 30 minutes, 6 – and/or 24 hours post amputation (MPA/HPA) using a MZ16F fluorescence stereomicroscope (Leica) combined with a ProgRes C3 camera (Jenoptik, Jena, Germany) or a Ts2-FL inverted microscope (Nikon) combined with a Ds-Fi3 colour camera (Nikon). For all pictures the exact same capturing settings were used. Additionally, all experiments were also performed without the carboxy-H2DCFDA-stain in order to discard possible autofluorescence at the wound sites.

### RNA Interference

Double-stranded RNA (dsRNA) for *Smed-egfr-3* and *Smed-egr-4* were synthesised as previously described (*Smed-egfr-3*, forward primer: GTACTGGGCAATGTTGGACCTGGC, reverse primer: TGACGGCCTCATGTGGGGATCATCG; *Smed-egr-4*, forward primer: GGCCGCGGTATGGGATATTCTTCTCAACTG: reverse primer: GTAATTATGAGTCGTGTAGGC). Animals were injected in two rounds of 3 consecutive days each with 4 days elapsed in between. On day 4 of the second round, planarians were amputated pre- and post-pharyngeally to induce regeneration. Injections were done using the Nanoject II (Drummond Scientific, Broomall, PA, USA) and consisted of three times 32 nl containing 1 µg/µl dsRNA. Controls were injected with dsRNA of *gfp*.

### In Situ Hybridization

Riboprobes for in situ hybridizations were synthesized using the DIG RNA labelling kit (Sp6/T7, Roche) following the manufacturer’s instructions. Primers used for the development of the DIG-labeled *Smed-MKP* probe were the following; forward primer: GACAATTTACGTTGTCCAACA, reverse primer: GTCCGGCGCCGTTTGACCCA.

To perform whole mount *in situ* hybridizations, regenerated trunk fragments were fixed at 6 hours post amputation (6HPA), using ice cold 2% HCl (ultrapure H_2_O, 5 minutes). Next, they were put in Carnoy’s solution (60% ethanol, 10% acetic acid and 30% chloroform) for 2 hours on 4°C. After 1 hour incubation in 100% MeOH at − 20 °C, samples were washed and bleached in 6% H_2_O_2_ (MeOH) for 16 h. Animals were rehydrated through a series of ethanol washes followed by PBSTween (10%) whashes. Afterwards, animals were treated with 20 µg/ml proteinase K (Ambion)/PBSTween for 10 minutes at RT. The proteinase K/PBSTween was removed with a PBSTween (5 minutes, 4°C) washing step and postfixed in 4% paraformaldehyde (PFA)/PBS for one hour at 4°C. Tissues were acetylated by 20 minutes of incubation in 0.1 mM TEA after which 25 µl of acetic anhydride was added. After 10 additional minutes another 25 µl of acetic anhydride was added. All the steps including TEA were performed at room temperature. Animals were washed with PBSTween (10 minutes, RT). Next, PBSTween was replaced by 50% prehybridisation buffer (in PBSTween) followed by incubation in 100% prehybridisation buffer (50% formamide, 5 × SSC, 0.1 mg/ml Yeast tRNA, 0.1 mg/ml heparina, 0.1% Tween 20, 10 mM DTT) for 2 hours at 56 °C. Hybridization was carried out for 16 hours at 56 °C in hybridization buffer (prehybridisation buffer + 10% Dextran Sulphate and the DIG-labelled probe). Afterwards, samples were washed through a series of posthybridisation buffers and buffer I ((0.1% Triton x100, Maleic acid 11.6 g, NaCl 9.76 g, 2N NaOH 95ml)/1000ml) and blocked during a 1h incubation in Buffer II (Buffer I with 10% Blocking Solution). Samples were incubated at RT for 3 hours in 1:2000 anti-DIG/Buffer II. Colour development was performed by incubation of the samples in 20 µl NBT/BCIP stock solution/ml TMN at RT. When the colour reaction reached the desired state, the animals were washed with PBS and fixed in 4% PFA/PBS. To optimize the colour development, a series of ethanol washes was performed. Finally, the samples were washed with PBS and kept in 70% glycerol/PBS at 4°C. Samples were analysed and bright-field images were digitized using the MZ16F stereomicroscope (Leica) equipped with a ProgRes C3 camera (Jenoptik, Jena, Germany).

## Data availability

The authors declare that all data supporting the findings of this study are available within the article and its Supplementary Information files or from the corresponding author upon reasonable request.

## Acknowledgements

The authors thank Yoshihiko Umesono for the anti-pERK antibody and Natascha Steffanie and Ria Vanderspikken for their skillful technical assistance. This work was financially supported by grants BFU2015-65704P and PGC2018-100747-B-100 to F.C. (Ministerio de Ciencia, Innovación y Universidades, Spain), and by FWO (1522015N, 1522719N and GOB8317N) and BOF UHasselt to K.S. The research leading to results presented in this publication was carried out with infrastructure funded by EMBRC Belgium - FWO project GOH3817N to K.S.

---

**Supplementary Figure 1.**
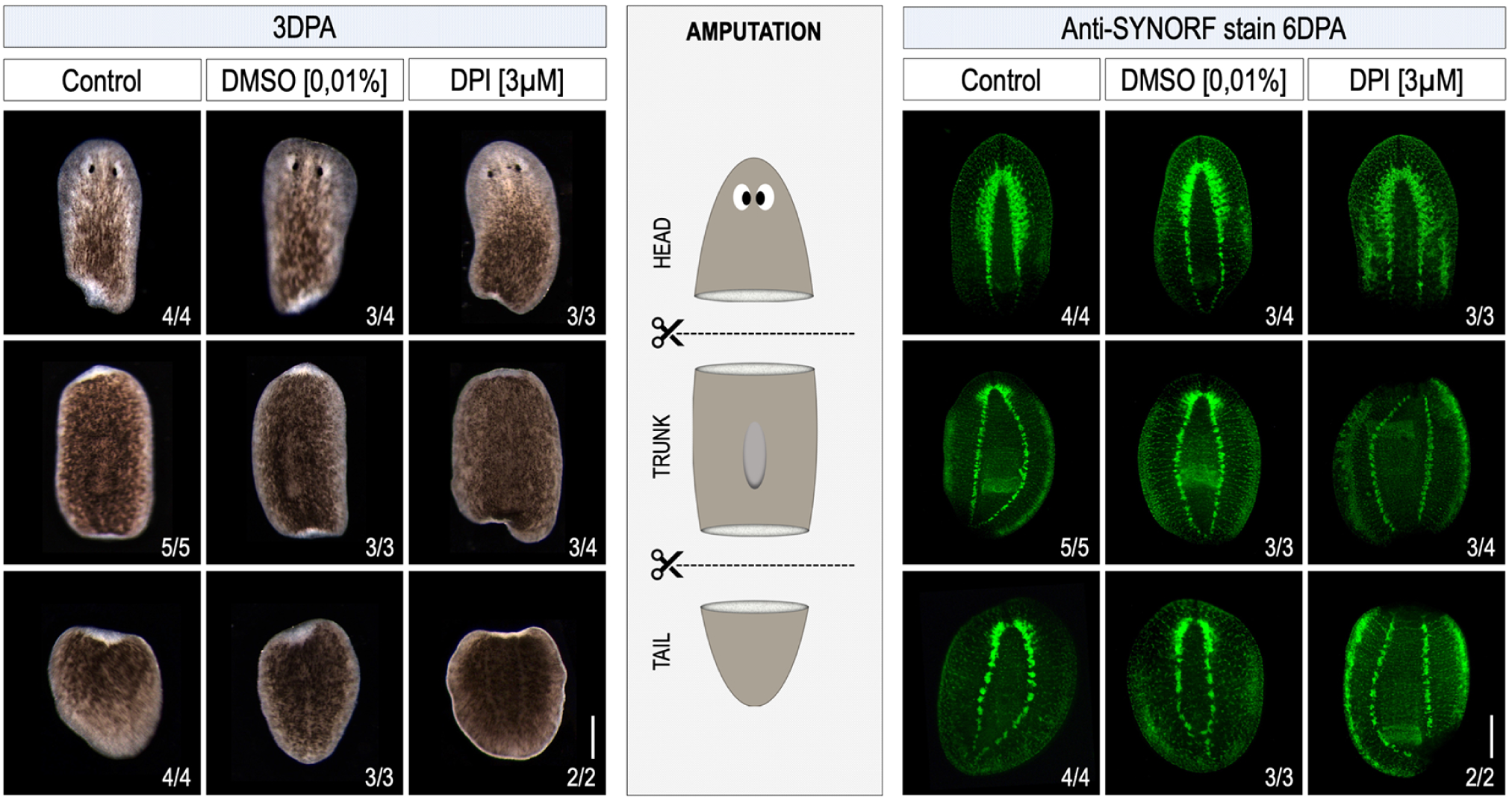
DPI mediated inhibition of ROS production impairs blastema formation and CNS regeneration. Control animals were kept in culture medium. In an additional control group, animals were exposed to 0,01% DMSO, while ROS inhibition was mediated by exposure to 3 mM DPI. In the left panels, all images were taken at 3 days post amputation (3DPA). The amputation setup is shown in the central panel. The right panels show regenerating fragments after a immunostaining with anti-SYNORF1 in order to visualise the central nervous system at 6 days post amputation (6DPA). Scale bar 200µm.

**Supplementary Figure 2.**
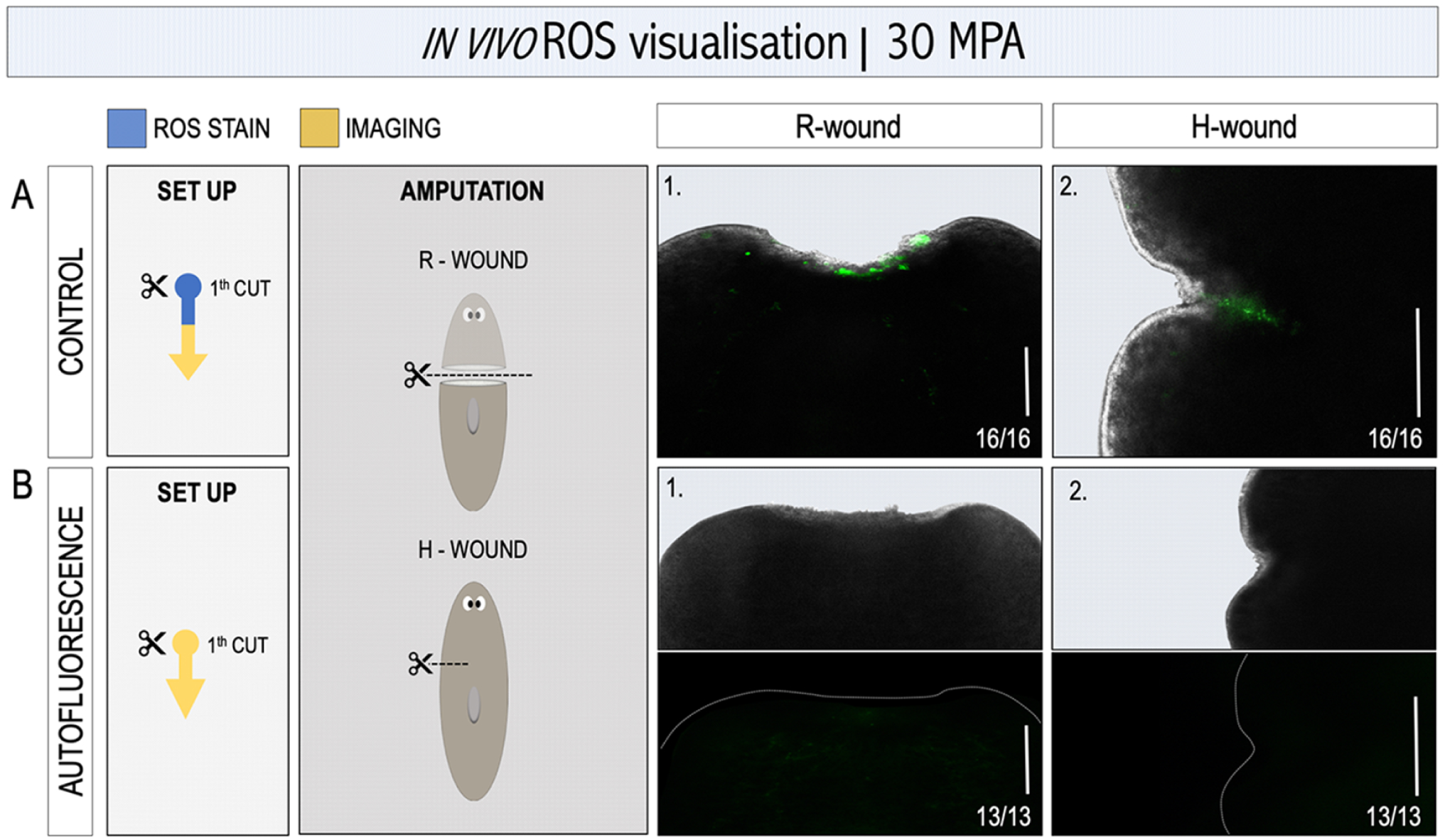
The fluorescent signal at the wound site is ROS production specific and not related to autofluorescence. For each condition, the experimental setup is displayed in the white box; blue: ROS visualisation procedure using carboxy-H_2_DCFDA, yellow: imaging procedure. Amputation setup is displayed in the light grey box. Regenerative wound (R-wound) & healing wound (H-wound) are indicated. All animals were visualised 30 minutes post amputation (MPA). A representative close-up image (merged or separated: bright field and fluorescence) of either an R-wound or H-wound is displayed on the right panel. (**A**) Control condition with ROS visualisation at the site of the R-wound (**A.1**) and H-wound (**A.2**). (**B**) No *in vivo* ROS stain is performed in order to exclude possible autofluorescence at the wound sites. In both R- (**B.1**) and H-wound (**B.2**), no autofluorescence was detected. Because of the absence of a fluorescence signal (**B**), the close-up of the wound site is shown in bright field (upper panel) and fluorescence (lower panel) separated. The dotted, white line indicates the border of the wound site. Scale bar 100µm.

**Supplementary Figure 3.**
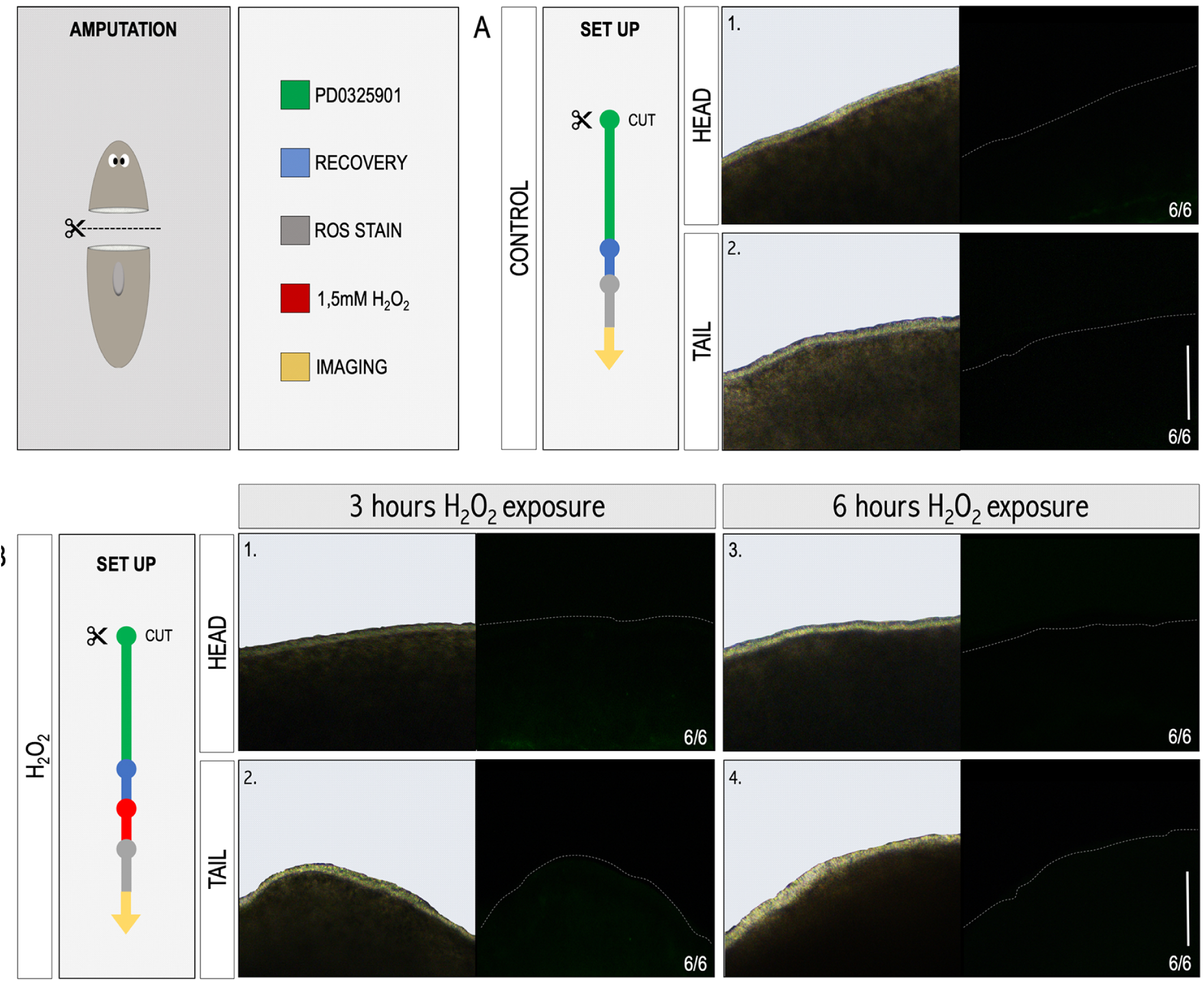
The rescue of regeneration in MEK-inhibited tails by H_2_O_2_ treatment is not attributed to rewounding by the exposure. For each condition, the experimental setup is displayed in the white boxes. Colour code as follows; green: MEK inhibition by PD0325901 (10 µM, 5 days), blue: recovery in fresh medium after several washes (1day), red: treatment with H_2_O_2_ (1,5 mM, 3 or 6 hours), grey: ROS visualisation procedure using carboxy-H_2_DCFDA, yellow: imaging procedure. The amputation setup is displayed in the grey box; amputation above the pharynx generating a head and tail fragment. H_2_O_2_ treated fragments were *in vivo* stained and imaged after either 3 - or 6 hours of exposure to H_2_O_2_. A close-up of the epidermis is presented in bright field (left panel) and fluorescence (right panel) separated, and is representable for the whole fragment. The dotted, white line indicates the border of the fragment. (**A**) control condition: MEK-inhibited fragments without any other intervention. (**B**) treatment conditions: MEK-inhibited fragments followed by 3-(**B.1/2**) or 6 hours (**B.3/4**) of H_2_O_2_-treatment. Scale bar 50µm.

